# A Pan-Respiratory Antiviral Chemotype Targeting a Host Multi-Protein Complex

**DOI:** 10.1101/2021.01.17.426875

**Authors:** Maya Michon, Andreas Müller-Schiffmann, Anuradha F. Lingappa, Shao Feng Yu, Li Du, Fred Deiter, Sean Broce, Suguna Mallesh, Jackelyn Crabtree, Usha F. Lingappa, Amanda Macieik, Lisa Müller, Philipp Niklas Ostermann, Marcel Andrée, Ortwin Adams, Heiner Schaal, Robert J. Hogan, Ralph A. Tripp, Umesh Appaiah, Sanjeev K. Anand, Thomas W. Campi, Michael J. Ford, Jonathan C. Reed, Jim Lin, Olayemi Akintunde, Kiel Copeland, Christine Nichols, Emma Petrouski, A. Raquel Moreira, I-ting Jiang, Nicholas DeYarman, Ian Brown, Sharon Lau, Ilana Segal, Danielle Goldsmith, Shi Hong, Vinod Asundi, Erica M. Briggs, Ngwe Sin Phyo, Markus Froehlich, Bruce Onisko, Kent Matlack, Debendranath Dey, Jaisri R. Lingappa, M. Dharma Prasad, Anatoliy Kitaygorodskyy, Dennis Solas, Homer Boushey, John Greenland, Satish Pillai, Michael K. Lo, Joel M. Montgomery, Christina F. Spiropoulou, Carsten Korth, Suganya Selvarajah, Kumar Paulvannan, Vishwanath R. Lingappa

**Affiliations:** Prosetta Biosciences, San Francisco, CA, USA; Institute of Neuropathology, Heinrich Heine University, Düsseldorf, Germany; Vitalant Research Institute, San Francisco, CA, USA; Veterans Administration Medical Center, San Francisco, CA, USA; University of Georgia, Animal Health Research Center, Athens, GA, USA; Institute of Virology, Heinrich Heine University, Düsseldorf, Germany; Santo Biotech, LLC., Pendleton, IN, USA; MS Bioworks, Ann Arbor, MI, USA; Dept. of Global Health, University of Washington, Seattle, WA, USA; Onipro LLC., Kensington, CA, USA; University of California, San Francisco, CA, USA; Viral Special Pathogens Branch, US Centers for Disease Control and Prevention, Atlanta, GA, USA

## Abstract

We present a novel small molecule antiviral chemotype that was identified by an unconventional cell-free protein synthesis and assembly-based phenotypic screen for modulation of viral capsid assembly. Activity of PAV-431, a representative compound from the series, has been validated against infectious virus in multiple cell culture models for all six families of viruses causing most respiratory disease in humans. In animals this chemotype has been demonstrated efficacious for Porcine Epidemic Diarrhea Virus (a coronavirus) and Respiratory Syncytial Virus (a paramyxovirus). PAV-431 is shown to bind to the protein 14-3-3, a known allosteric modulator. However, it only appears to target the small subset of 14-3-3 which is present in a dynamic multi-protein complex whose components include proteins implicated in viral lifecycles and in innate immunity. The composition of this target multi-protein complex appears to be modified upon viral infection and largely restored by PAV-431 treatment. Our findings suggest a new paradigm for understanding, and drugging, the host-virus interface, which leads to a new clinical therapeutic strategy for treatment of respiratory viral disease.

## Background

The current SARS-CoV-2 pandemic has been characterized by waves of infection. Emerging mutants, with varying degrees of resistance to current vaccines and waning immune responses within the population, have contributed to the seemingly-unending surges of disease (1,2). Furthermore, the risk of a new pandemic, from avian influenza, respiratory syncytial virus (RSV), or another virulent pathogen known to exist in animal reservoirs, is ever present (3). Given how rapidly SARS-CoV-2 spread across the globe once it had been transmitted to humans, concern about highly pathogenic respiratory viruses should not be considered as an abstract, hypothetical threat (4). A technical solution is needed which can account for the degrees of uncertainty and variation inherent to pandemic preparedness and response efforts. Otherwise, antiviral countermeasures will continue to aim at an ever-moving target and always be one step behind. In this paper we will propose a novel solution—one small molecule compound with potent activity against all six families of viruses that cause most respiratory viral disease in humans.

Viruses in *Adenoviridae*, *Coronaviridae*, *Herpesviridae*, *Orthomyxoviridae*, *Paramyxoviridae*, and *Picornaviridae* families cause over 95% of respiratory disease in humans (5). Diversity between these viral families, which include both DNA and RNA viruses, and viruses that are both enveloped and not, is extremely broad (5). The drugs which are available to treat some of these viruses target the varying proteins encoded by the different viral genomes (6–8). Oseltamivir (Tamiflu) and zanamivir (Relenza) work on influenza by inhibiting neuraminidase, a viral enzyme that propagates infection by facilitating the spread of viral particles throughout the host (8). Acyclovir, a treatment for herpes simplex virus, inhibits viral DNA polymerase (6). Paxlovid, the new drug for SARS-CoV-2, is a protease inhibitor that blocks viral enzymes responsible for catalyzing critical maturation steps within the virus’s lifecycle (7). But since any one of these viral families represents a small minority of respiratory viral cases, a diagnosis must be made before potentially effective treatment is initiated. Yet considerable evidence suggests that the earlier the treatment, the greater is the efficacy (9).

Host-targeted antiviral drugs have been proposed as a new strategy for antiviral drug development (10–14). Viruses can only reproduce successfully if they are able to redirect host machinery to suit viral needs (e.g. by building its capsid, blocking immune response, etc.) rather than the needs of the host, which is to maintain homeostasis (15). The viral generation time is several orders of magnitude shorter than the host’s, making it likely that the host-virus interactome has been highly selected by viral evolution to provide the best way to reprogram host machinery (16,17). While viruses employ a range of strategies for hijacking host machinery, “high value” sites of host-viral interface are likely to be exploited by more than one family of virus. Those sites would make ideal targets for pan-family antiviral drugs, but identifying them is a challenge.

We hypothesized that it would be possible to identify these high-value host-viral interface sites, and develop drugs which target them, using cell free protein synthesis and assembly (CFPSA) systems (13,18,19). Cell free systems have been used to observe and understand critical molecular-level processes since 1897 when Eduard Buchner demonstrated that cell-free extracts could carry out the same fermentation reactions as living cells (20). More recently, cell-free protein synthesis has been a critical tool used to decipher the genetic code, deconvolute protein trafficking, and functionally reconstitute the transient virus-host-protein interactions that culminate in viral capsid formation (21–25). The last of these applications, which gave rise to the observation that viral capsid assembly in the cell-free system is dependent on both host machinery and metabolic energy, and thus cannot be due to spontaneous self-assembly, provided the rationale for developing our antiviral drug screen. Our hypothesis was that if viral capsid assembly is a host-catalyzed process, then antiviral therapeutics could be developed by inhibiting the critical host enzymes co-opted by a virus to catalyze assembly of its capsid. To test this hypothesis, we set up a phenotypic screen for compounds that could block viral capsid formation in the CFPSA system, without inhibiting protein synthesis (13,19).

There are several advantages of a CFPSA-based drug screen. First, it uniquely serves to magnify early events in protein biogenesis that would otherwise be obscured by events in the rest of a protein’s life within the cell. Second, it recreates the reality of protein heterogeneity, including with respect to post-translational modifications (PTMs, (26–28) and multi-protein complex formation (29–31). Finally, it exploits the recent appreciation that critical events in protein-protein interactions may occur co-translationally, that is, while a protein is nascent (32–36). While in principle such a screen could detect direct binders of the translated viral protein(s), we suspected that the effect of binding a catalytic host target would be much greater, since blocking one enzyme affects many substrate molecules and in this case, the viral capsid monomer would effectively be the substrate for catalyzed capsid assembly.

There is an presumption that drugs which target host proteins pose an inherent risk of toxicity (14). However, one implication of the burgeoning literature in favor of “moonlighting” functions of proteins is that only a small subset of any given protein participates in any particular MPC (37–39). Once a hit compound was identified by the CFPSA screen it should then be possible to drive its structure-activity relationship to selectivity for the relevant subset of the target protein. We therefore anticipated the need to defer full assessment of toxicity until after structure-activity relationship (SAR) advancement of initial hits. Thus, once an antiviral compound targeting the host were identified by CFPSA, it could subsequently be advanced, first for efficacy, and then to moderate toxicity. This could be achieved either by virtue of the target being a small subset of the full complement of that protein in the cell, or if the virus modified the host target for its needs, SAR might be selectively tuned to the form of the target needed by the virus.

The results, to be provided in this paper, focus on the advancement of one novel chemical series identified as a viral assembly modulator in the CFPSA screen, that appears to show pan-family antiviral efficacy in cells and animals. Experiments were performed to advance the potency of this antiviral chemical series and better understand its target and mechanism of action, to provide an understanding of this new host-viral interface.

## Results

### Identification and assessment of early assembly-modulating hit compounds PAV-773 and PAV-835

A cell-free protein synthesis and assembly (CFPSA) based phenotypic screen was established for influenza (FLUV) analogous to what has been done for rabies, HIV, and other viruses (13,19,40,41). Unlike conventional phenotypic screens, this screen was carried out in cellular extracts rather than in living cells. The phenotype being screened was the ability of newly synthesized viral capsid protein for form multimers. In the CFPSA system, faithful formation of multimeric capsid protein complexes is a quantifiable, functional endpoint (see diagram in **Figure 1A**).

**Figure 1.**
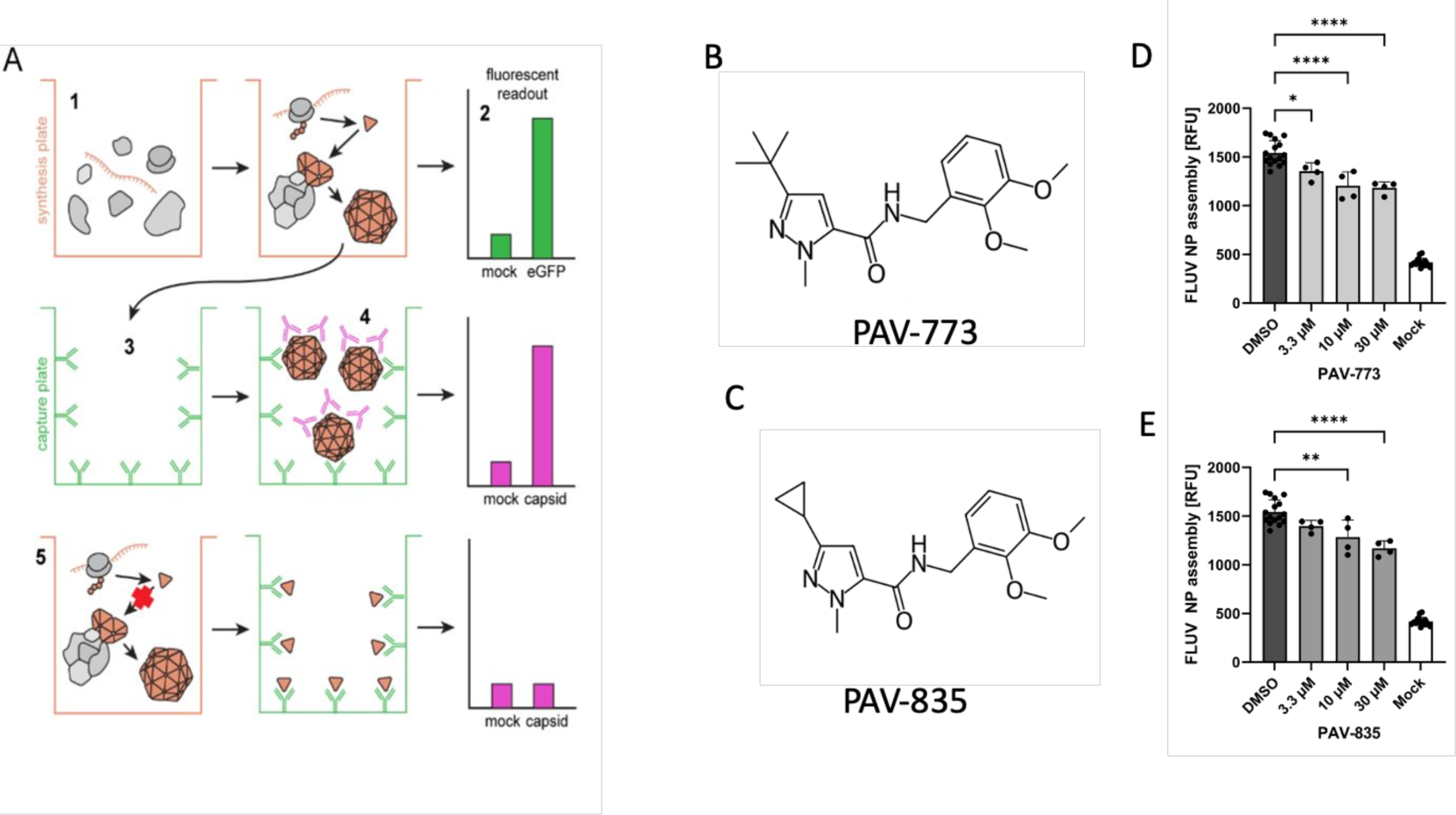
Identification of PAV-773 and PAV-835 as FLUV assembly inhibitors. **Figure 1A** shows a schematic of the CFPSA phenotypic drug screen indicating steps and readouts. CFPSA reactions carried out in a 384 well plate format **(1)** result in synthesis of encoded FLUV proteins, with co-expression of eGFP to distinguish compounds that lower fluorescence readout due to a trivial effect on a protein synthesis **(2)**. Assembled products are transferred to a capture plate **(3)** which is coated with antibodies to the FLUV nucleoprotein, capturing and immobilizing synthesized FLUV proteins. As a function of multimerization, unoccupied epitopes will exist on bound assembly Intermediates and completed viral structures. Secondary antibodies with fluorescent tags bind those exposed epitopes **(4)**, generating a fluorescent readout specific to multimeric assembly. Drug action that directly or indirectly blocks multimer formation results in a diminution of signal **(5)**. **Figure 1B** shows the chemical structure of PAV-773 and **Figure 1C** shows the chemical structure of PAV-835, early hits in the CFPSA screen. **Figure 1D** and **Figure 1E** show the effects of PAV-773 and PAV-835 respectively at 3.3uM, 10uM, and 30uM doses on assembly of FLUV NP in the screen, compared to DMSO and a mock negative control. Average relative fluorescent units (RFU) detected from quadruplicate-repeat samples are graphed with standard deviation shown as error bars and statistical significance calculated on GraphPad Prism using an ordinary one-way ANOVA test is indicated by asterisks.

From a library of 150,000 drug-like small molecules, 30,400 compounds were screened and compounds that interfere with the biochemical pathways of host-catalyzed FLUV capsid assembly were identified as hits. PAV-773 and PAV-835 were early compounds from a chemical series identified in the screen as inhibitors of FLUV capsid assembly (see **Figures 1B** and **1C** for their respective chemical structures). Both compounds blocked assembly of FLUV nucleoprotein into a completed capsid in a dose-dependent manner, relative to control (see **Figures 1D** and **1E** for their respective activity against FLUV capsid assembly).

The FLUV antiviral activity of PAV-773 and PAV-835 was validated against infectious virus in MDCK cells by TCID_50_ determination (see **Figures 2A**). The effective concentration for half maximal activity (EC50) against infectious FLUV for both PAV-773 and PAV-835 were lower than 1uM (see **Figure 2A**).

**Figure 2.**
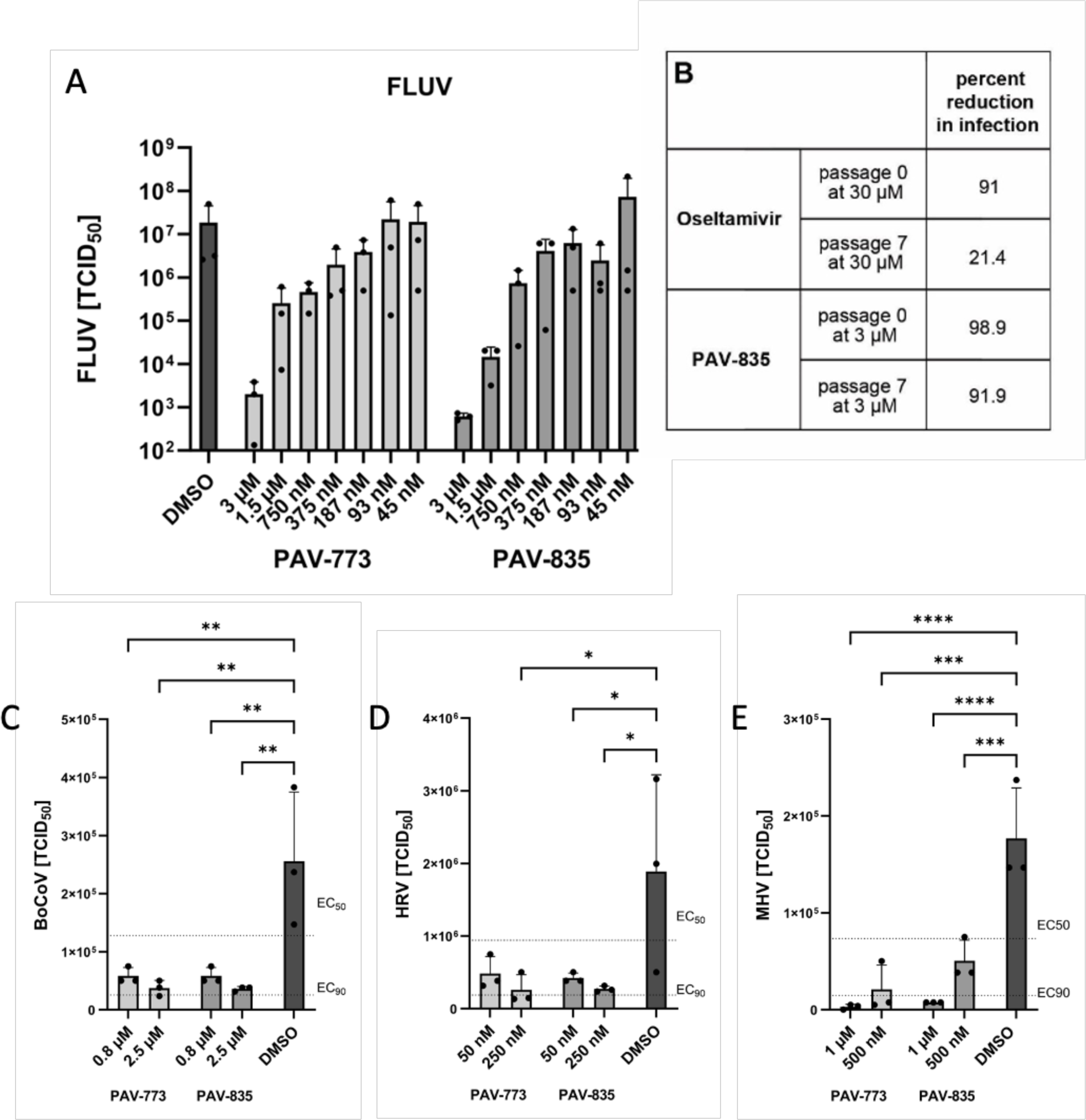
Validation of PAV-773 and PAV-835 antiviral activity in cell culture and evidence for a barrier to resistance development. **Figure 2A** shows activity of PAV-773 and PAV-835 against infectious FLUV (A/WSN/33) in MDCK cells by TCID_50_ determination. Averages and standard error of triplicate samples are graphed, and statistical significance calculated by one-way ANOVA on GraphPad Prism is indicated with asterisks. **Figure 2B** shows activity of PAV-835 or oseltamivir against FLUV (A/WSN/33) in MDCK cells after 7 passages in the presence of compound. **Figure 2C** shows activity of PAV-773 and PAV-835 against infectious coronavirus (BRCV-OK-0514-2) in HRT-I8G cells by TCID_50_ determination. **Figure 2D** shows activity of PAV-773 and PAV-835 against infectious rhinovirus (HRV-16) in HI-HeLa cells by TCID_50_ determination. **Figure 2E** shows activity of PAV-773 and PAV-835 against infectious herpesvirus (MHV-68) in BHK-21 cells by TCID_50_ determination. PAV-773 and PAV-835 both displayed EC50s of less than 1uM for all four viruses studied. Statistical significance for **Figures 2C-E** were calculated by ordinary one-way ANOVA tests on GraphPad Prism and are indicated with asterisks.

The emergence of viral resistance is a common challenge for the development of effective antiviral therapeutics (42). Oseltamivir (Tamiflu), an antiviral small molecule targeting FLUV neuraminidase, is known to select for viral resistance mutants (43). To assess the propensity for FLUV to gain resistance to our chemotype, MDCK cells were infected with serial passages of FLUV in the presence of PAV-835. With each passage, the infected media was used to infect fresh MDCK cells. Higher concentrations of compound were added with each passage to drive resistance (93.5nM to 3uM). After 7 passages with compound, PAV-835 retained the same activity against FLUV as it did against a naive strain which had been passaged for 7 times without compound, demonstrating a barrier to the development of resistance (see **Figure 2B**). In parallel, the same experiment was conducted using Oseltamivir (ranging from 935nM to 30uM), antiviral resistance developed and the compound lost activity by passage 7 (see **Figure 2B**).

We counter-screened PAV-773 and PAV-835 for activity against other viral families by assessment of viral titer in cell culture by TCID_50_ (see **Figures 2C-E**). Both compounds were found to have EC50s of less than 1uM against bovine coronavirus (BoCoV), human rhinovirus (HRV), and murine herpesvirus (MHV). These data led us to refer to these compounds as *<u>pan-respiratory viral assembly modulators</u>* based on their initial identification as modulators of FLUV capsid assembly and subsequent demonstration of efficacy against multiple respiratory-disease causing viruses.

### Validation of the antiviral activity of PAV-773 and PAV-835 in animals

At the time we were characterizing the activity of these early compounds, an outbreak of Porcine epidemic diarrhea virus (PEDV) led to the loss of more than 10% of the pig population in the United States (44). Since PEDV is a member of the coronavirus family, we predicted that while the chemical series was early in the drug-development process, the compounds would likely show antiviral activity against PEDV. PAV-773 and PAV-835 were assessed in outbred pigs randomized within each litter into control and treatment groups and infected with PEDV. Both compounds significantly increased likelihood of survival, relative to the control (see **Figures 3A** and **3B**).

**Figure 3.**
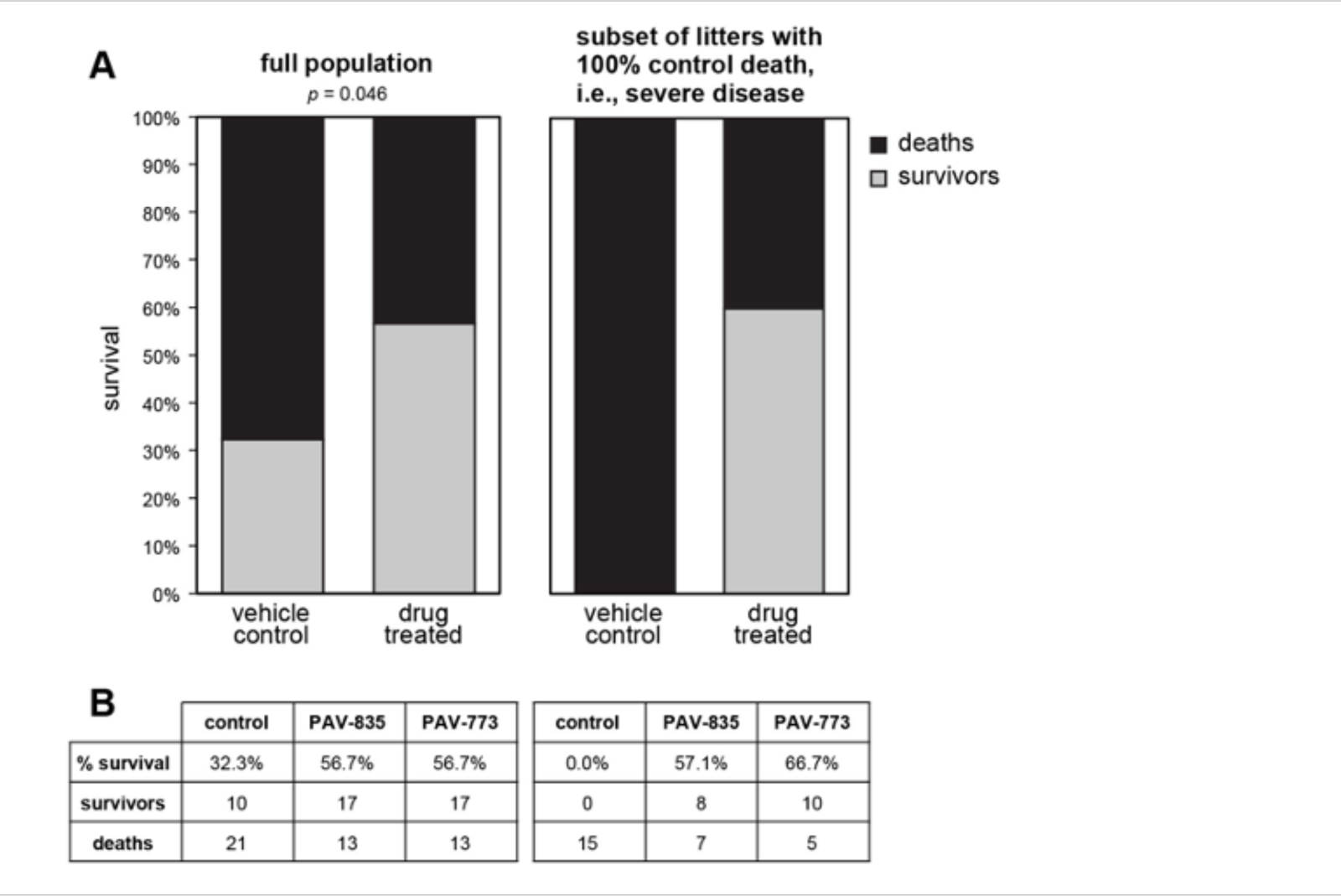
Validation of PAV-773 and PAV-835 antiviral activity in pigs. Pigs were randomized into control and treatment groups then infected with PEDV, a pig coronavirus. **Figure 3A** Left is shown the percent survival for all animals in the study. The p-value was calculated on GraphPad Prism using Fisher’s exact test. Right is shown the percent survival in the subset of litters in which all animals in the randomized control (vehicle) treatment group died. As can be seen, compound treatment is equally efficacious in this severe disease subset. **Figure 3B** shows the breakdown of survival for PAV-773 and PAV-835.

### Characterizing the antiviral activity of PAV-431, a more advanced analog from the Pan-Respiratory Assembly Modulator chemical series

A structure-activity-relationship (SAR) was pursued to advance the pan-respiratory assembly modulator chemical series emerging from the early hits, and to understand how changes to the chemical structure altered activity against infectious FLUV (see **Supplemental Figure 1**). PAV-431 was identified as a chemical analog with improved efficacy (see **Figure 4A** for its chemical structure and **Supplemental Figure 2A** for its synthetic scheme).

**Figure 4.**
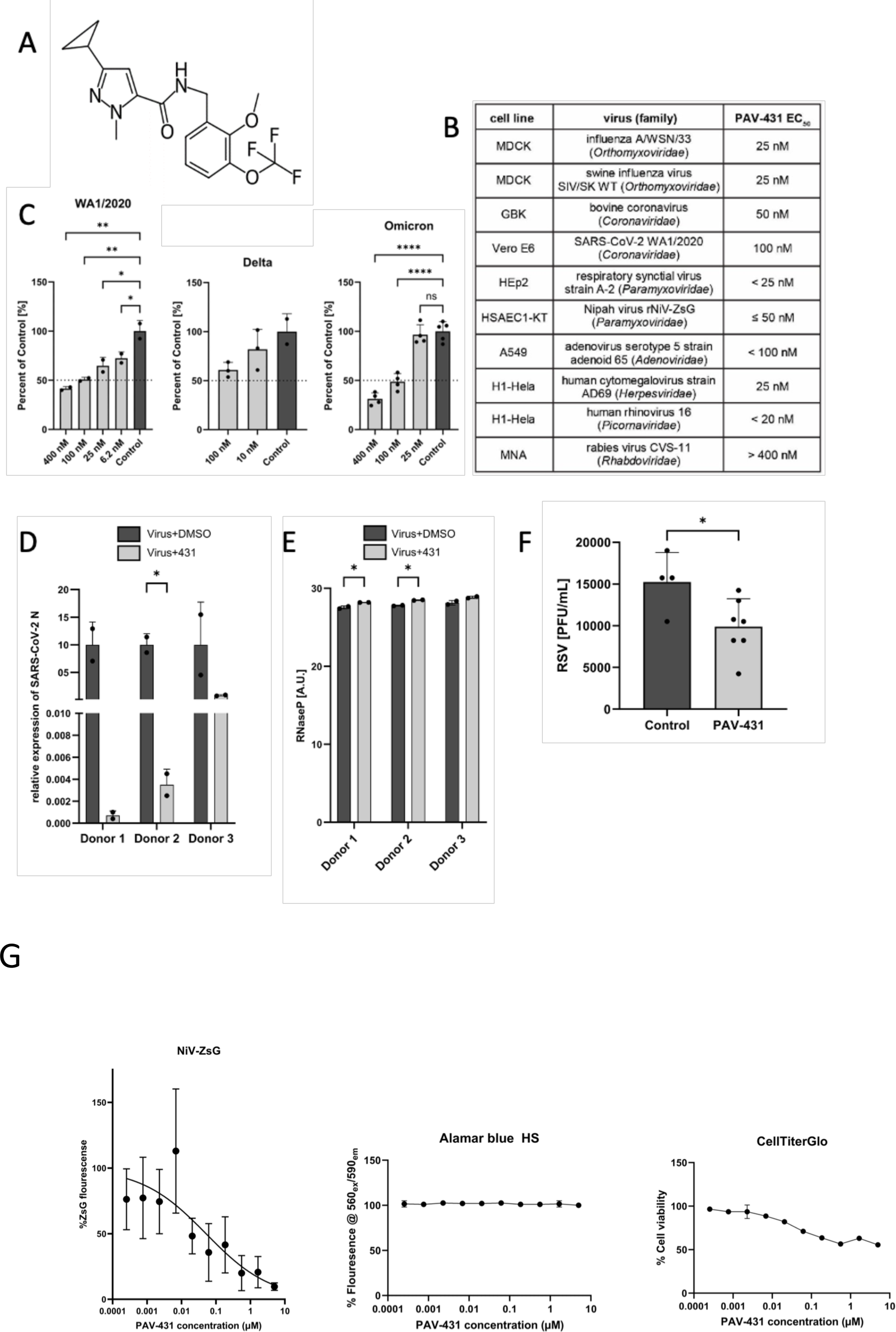
Antiviral activity of PAV-431 against all viral families which cause respiratory disease in humans. **Figure 4A** shows the chemical structure of PAV-431, an analog from the pan-respiratory assembly modulator chemical series. **Figure 4B** shows the efficacy of PAV-431 against multiple viruses in cell culture by TCID_50_ where and EC50 of 100nM or lower is observed for every family of virus causing human respiratory disease. **Figure 4C** shows dose dependent antiviral activity of PAV-431 compared to a DMSO control against multiple SARS-CoV-2 strains: (Wa/2020, lineage A) in Vero E6 cells, determined by plaque assay, delta variant (lineage B.1.617.2) and omicron variant (lineage B.A.1) in Calu-3 cells determined by qPCR of the SARS-CoV-2 N gene and/or TCID_50_. Data shown are the averages of three biological replicates where error bars indicate standard error. Statistical significance was calculated on GraphPad Prism using an ordinary one-way ANOVA test for each dataset. **Figures 4D** and **4E** show efficacy and nontoxicity of PAV-431 in primary human airway epithelial cells at air-liquid interface. Bronchial epithelial cells from three lung donors were culture to an air-liquid interface, infected with SARS-CoV-2 (gamma variant, lineage P.1) and treated with either PAV-431 or vehicle. **Figure 4D** shows average of two replicates with error bars indicating standard error where viral replication was determined by qPCR measurement of the SARS-CoV-2 N gene. **Figure 4E** shows a lack of observed toxicity assessed by levels of RNase P. **Figure 4F** shows results of PAV-431 in an animal efficacy trial against RSV in cotton rats. Averages are shown where error bars indicate standard error. A significant drop in viral titer was observed with PAV-431 treatment, relative to vehicle (unpaired t-test p=0.016). The statistical significance for 4D-F was calculated on GraphPad Prism using unpaired t-tests. **Figure 4G** shows the efficacy of PAV-431 against Nipah virus, a BSL-4 member of the *Pneumoviridae* family in human telomerase reverse-transcriptase immortalized primary-like small airway epithelial cells (HSAEC1-KT, ATCC CRL-4050) cultured in Airway Epithelial Basal Medium (ATCC) supplemented with Bronchial Epithelial Cell Growth Kit (ATCC). For infections and cell viability assays (done in duplicate) by both Alamar Blue and Cell Titer glo, HSAEC1-KT cells were cultured with growth medium with 5 mM of D-glucose solution (Gibco). PAV-431 was added prior to infection as described in methods. As shown, CC_50_ of PAV-431 by Alamar Blue is > 10uM and by Cell titer glo is > 5uM, under these conditions.

PAV-431 was assessed by TCID50 for activity against multiple viral families. PAV-431 displayed an EC50 between 25nM and 100nM (depending on the virus) against members of *Orthomyxoviridae*, *Coronaviridae*, *Paramyxoviridae*, *Adenoviridae*, *Herpesviridae*, and *Picornaviridae*—all six families of viruses which cause respiratory disease in humans (see **Figure 4B**). Within *Coronaviridae*, PAV-431 showed efficacy against the WA 1/2020, delta, and omicron strains of SARS CoV-2 (see **Figure 4C**). Finally, PAV-431 was assessed against Nipah virus, a BSL-4 member of the *Pneumoviridae* family with pandemic potential should it ever jump species and become capable of human-to-human aerosol transmission and shown comparably potent (see **Supplementary Figure 3**).

In addition to demonstrating efficacy in transformed cells, PAV-431 showed activity against the gamma variant of SARS CoV-2 in primary human bronchial epithelial cells cultured to an air-liquid interface (ALI) (see **Figure 4D**). 100nM PAV-431 eliminated approximately 90% of more of viral load compared to vehicle treatment in three ALI studies derived from three different human lung donors without inflicting significant toxicity to the cells, as measured by levels of RNASe P (see **Figure 4D** and **4E**). Notably, PAV-431 did not show significant activity at those doses against rabies virus, indicating that even though the compound displays broad pan-family efficacy, there is some selectivity for a target present in some, but not all, viral families (see **Figure 4B**).

We assessed the degree to which the chemical properties of PAV-431 meet the standard criteria for advancement into a drug candidate. PAV-431 displayed promising properties including being negative for hERG channel inhibition, and without substantial Cerep panel enzyme inhibition (see **Supplemental Figure 3**). When administered to rats, a dose of 5mg/kg administered intraperitoneally (IP) was found to be safe, reaching a concentration of 293 ng/ml in plasma and 452 ng/nl in lungs (see **Supplemental Figure 3A**).

Given the respectable PK properties, PAV-431 was tested in cotton rats infected with respiratory syncytial virus (RSV), a paramyxovirus, to assess animal efficacy for a more advanced compound in the series against a second family of respiratory disease-causing viruses. A small but statistically significant drop in RSV titer was observed with PAV-431 treatment, relative to the vehicle-only control (see **Figure 4F**).

### Identifying the molecular target of the Pan-Respiratory Assembly Modulators

Since the pan-respiratory viral assembly modulator chemical series had been validated as significantly active in cellular or animal models for seven viral families, we sought to understand the molecular target being acted upon by the compound in order to achieve the results. To identify the target, PAV-431 was coupled to an Affi-gel resin from a position on the molecule unrelated to the activity based on SAR exploration. Once bound to a resin, it could serve as a target-binding ligand for drug resin affinity chromatography (DRAC) (see **Supplemental Figure 2B** for synthetic scheme of a PAV-431 resin).

In the DRAC protocol, extracts were prepared from MRC-5 cells which were uninfected, infected with either FLUV or BoCoV, and treated with 400nM PAV-431 or an equivalent amount of DMSO. The extracts were applied to the PAV-431 resin or a control resin containing an Affi-gel matrix bound to itself, washed with 100 bed volumes of buffer, eluted with 100uM PAV-431, then stripped with 1% SDS. In the course of these studies, we discovered that providing metabolic energy substrates in the form of nucleotide triphosphates greatly enhanced target formation (45). We established conditions for energy-dependent drug resin affinity chromatography (eDRAC) by supplementing the extract and the elution buffer with a “energy cocktail” of ribonucleotide triphosphates 1mM rATP, 1mM rGTP, 1mM rCTP, 1mM UTP) and 5 ug/mL creatine kinase, with the binding, washing, and eluting steps conducted at 22°C.

When eDRAC eluates from the PAV-431 resins and the control resin were collected and analyzed by silver stain compared to the starting extract, several striking observations were made. While the protein profile in the starting extracts for uninfected, FLUV, and BoCoV infected cells appeared similar, the PAV-431 resin eluates were strikingly different, with a protein pattern not observed for free drug eluates from control resin (lacking the drug as an affinity ligand, see **Figure 5A**).

**Figure 5.**
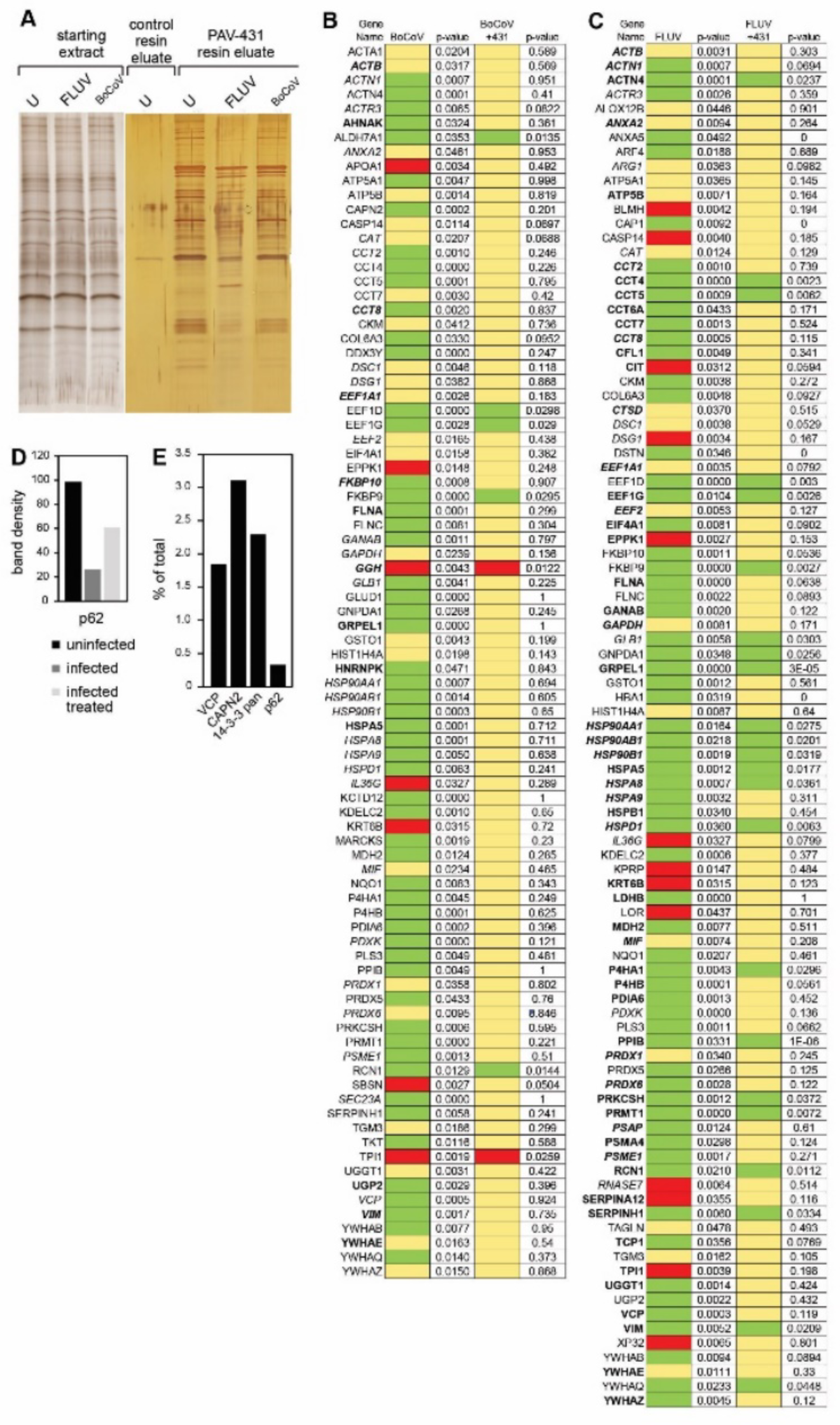
Protein composition of the PAV-431 eluate. eDRAC experiments were performed where uninfected, infected, or infected/PAV-431 treated cellular extract was incubated on a resin coupled to either PAV-431 or a 4% agarose matrix lacking the covalently bound drug. **Figure 5A** shows silver stain of a SDS-PAGE gel comparing protein composition of the starting cellular extract and the PAV-431 eluate for uninfected, FLUV infected and BoCOV infected MRC5 cells. **Figure 5B** shows MSMS analysis indicating protein composition and comparing log2 fold change and p-values in protein in triplicate repeated uninfected, FLUV infected, and FLUV/PAV-431 treated conditions. **Figure 5C** shows MSMS analysis indicating protein composition and comparing log 2 fold change in protein in triplicate-repeated uninfected, BoCoV infected, and BoCoV infected/PAV-431 treated conditions. Green indicates log2 fold change >1. Yellow indicates log2fold change between −1 and 1 (no change). Red indicates log2 fold change >-1. P values indicate significance of the findings. Where the gene product has been listed in bold font, indicates the protein is implicated in the literature as part of the host-virus interactome. Where the gene product has been listed in italic font, indicates the protein is implicated in the literature as related to innate immune system function. **Figure 5D** shows quantitation of the protein band detected by western blot analysis of the uninfected, infected, and FLUV infected/PAV-431 treated eluates for the protein p62/SQSTM1. **Figure 5E** shows quantitation of the protein band detected by western blot analysis of eDRAC from pig lung extract where starting extract and eluate were compared side-by-side and the amount of protein in the eluate is graphed as a percentage of the total amount of that protein present in the cell extract. Approximately 2% of the cellular VCP, 3% of the cellular CAPN2, 2.5% of the cellular 14-3-3 and 0.5% of the cellular p62 was found in the PAV-431 eluate.

Triplicate-repeat samples of eDRAC eluates generated from MRC-5 cell extract were sent for analysis by tandem mass spectrometry (MS-MS) to determine their protein composition. To analyze the data, LFQ intensity values for proteins identified in each condition were measured and compared against each other to generate log2 fold change values for each protein and each combination of conditions to provide a clear description of the differences observed under treatment conditions. Of 64 proteins identified by LFQ as increased in eluates upon FLUV infection, 41 are restored to uninfected levels after treatment with PAV-431 (see **Figure 5B**). All 13 proteins lost from eluates upon FLUV infection are restored to uninfected levels after treatment with PAV-431 (see **Figure 5B**). Of 56 proteins found increased in eluates upon BoCoV infection, 51 are restored to uninfected levels after treatment with PAV-431 (See **Figure 5C**). Of 7 proteins lost from eluates with BoCoV infection, 5 are restored to the uninfected levels after treatment with PAV-431 (See **Figure 5C**).

Proteins found to be significantly enriched or depleted by infection and/or treatment were searched in databases for known virus-host interactions and implication in the innate immune system interactome and many such proteins were identified (See **Figures 5B-5D**) (46–51). P62/SQSTM1, a regulator of innate immunity, was identified in the PAV-431 eluate by western blot. As with changes in protein composition observed by MS-MS, the amount of P62 decreased with FLUV infection but was restored with PAV-431 treatment (See **Figure 5D**).

The eDRAC protocol was also conducted with extract prepared from uninfected pig lung homogenate, rather than MRC-5 cells, and samples were analyzed by western blot. When analyzed side-by-side with an aliquot of the total starting material, it was determined that for particular proteins found in the eluate including VCP, CAPN2, 14-3-3, and P62, only a single digit percent, or less, of the total amount of specific proteins present in the extract was found in the PAV-431 eluate (See **Figure 5E**). The large majority of the component proteins did not bind to the resin, or bound nonspecifically such that they were removed with washing, with no significant further binding of drug resin flowthrough applied to a second copy of the drug resin.

To determine the relationship the proteins identified in the eluate had to one another, and to the compound, the eDRAC protocol was modified for photocrosslinking. An analog of PAV-431 was synthesized with diazirine and biotin moieties added to the same position at which the resin had previously been attached (See **Supplemental Figure 2C** for synthetic scheme and chemical structure of photocrosslinker analog). The photocrosslinker analogs were designed so that after an incubation with cell extract that would allow the compound to bind its target, exposure to ultraviolet light would form a covalent bond between the diazirine moiety of the compound and the nearest protein neighbor (52). The sample could then be solubilized and precipitated with streptavidin beads (which bind biotin with extremely high affinity) to identify the covalently crosslinked drug-binding proteins. The streptavidin precipitation (SAP) could be done using a native sample, which would pick up the direct drug binding protein(s) and with it, co-associated proteins that were part of an MPC. Alternatively, the SAP could be done using a crosslinked sample that was then denatured by treatment with SDS to 1% and DTT to 1mM with heating to 100oC for 3 minutes to denature all proteins, after which excess 1% Triton-X-100 buffer was added to take free SDS into Triton micelles. Use of this material for SAP would, by virtue of the covalent bond to the biotin containing diazirine-drug conjugate, identify only the direct drug-binding protein(s), with all other associated proteins lost upon denaturation and washing.

Uninfected pig lung was incubated on the PAV-431 resin under eDRAC conditions, washed 100x, eluted with the PAV-431 crosslinker analog, then exposed to ultraviolet light. The samples were then divided into two equal parts where one was left native and the other denatured, then both were adjusted to non-denaturing conditions and incubated with streptavidin beads. Blots of the SAP samples for VCP, CAPN2, and P62 showed those proteins in the native but not denatured samples, indicating that they were non-covalently co-associated with the compound, and therefore were not its direct binding partner (see **Figures 6A, B**, and **D**). Blots of the SAP samples for 14-3-3 showed nearly equal amounts of protein in both the native and denatured conditions, indicating that PAV-431 directly binds to 14-3-3 (See **Figure 6C**).

**Figure 6.**
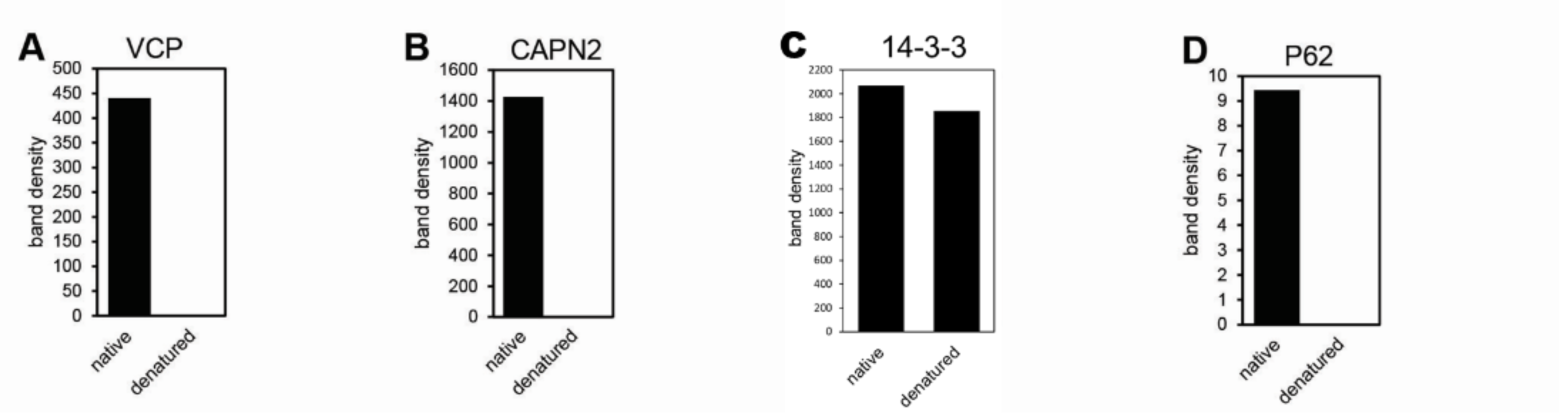
A cellular sub-fraction of 14-3-3 as the direct binding partner in an MPC drug target. eDRAC was conducted with pig lung homogenate extract eluted from the PAV-431 resin with the PAV-431 crosslinker analog. Eluates were exposed to UV light and precipitated with streptavadin in native and denaturing conditions then analyzed by western blot. **Figures 6A-D** shows quantitation of the protein for VCP, CAPN2, 14-3-3, and P62.

## Discussion

The antiviral chemotype studied here exhibits several notable features. These include activity across a broad range of respiratory viral families, a demonstrated barrier to development of viral drug resistance, different forms of the target present in uninfected vs infected cells, and substantial restoration of the target to the uninfected form with drug treatment. In all cases, a subset of the host protein 14-3-3 appears to be the direct drug-binding protein and is present within a large multiprotein complex notable for its transience and energy-dependence. PAV-431 binds both forms of the target, i.e. that present in uninfected cells and that present in infected cells, roughly equally well. Elsewhere a more advanced compound will be described that appears to be selective for the form present only in infected cells which includes, for SARS-CoV-2, the viral nucleoprotein, which is subsequently lost from the target upon drug treatment (53, 54).

Two general approaches can be taken for discovery of chemical compounds with therapeutic potential— target-based and phenotypic methods (55). Target-based methods of drug discovery involve screens that measure a small molecule’s interaction with a particular disease-implicated protein. Phenotypic methods involve screens that monitor how small molecules affect particular biochemical or physiologic readouts within model systems without requiring any prior knowledge of the protein target. It has recently been observed that most drugs have been discovered by variations on phenotypic screening (56). Most phenotypic screens involve whole cell assays (57). Such screens, while often successful, face significant drawbacks. The presence of confounding events in the complex milieu of a living cell can mask detection of potentially interesting targets. Moreover, feedback effects are typically complex and multifaceted (58–63). This can create a signal-to-noise problem for detection of potential contributors to a particular phenotypic effect. If multiple contributors are involved in creating a phenotype it may be hard to de-convolute the relationship of any given one to the behavior of the compound.

By contrast, the CFPSA-based phenotypic screening approach taken here focuses attention on those events set into motion early in protein biogenesis (during and immediately after protein synthesis). This results in an improved signal-to-noise ration by excluding much of the rest of the lifecycle of most proteins, for both viruses and cells, as confounding variables. A growing literature supports the notion that protein assembly is co-translational (32,64). Thus, CFPSA reveals aspects of the viral lifecycle not easily discernable by other methods.

14-3-3, the protein identified as PAV-431’s direct target (see **Figure 6**) is known to regulate multiple signaling pathways, including cell cycle progression, apoptosis, autophagy, and glucose metabolism, through protein-protein interactions (65–69). However, it has been difficult to convert these insights on 14-3-3 biology into therapeutic successes, perhaps because of this “promiscuity” of 14-3-3 (67–75).

The 14-3-3 targeting antiviral chemotype identified through CFPSA is promising precisely because it does *not* target all of 14-3-3, but rather a tiny subset found within a particular transient, energy-dependent MPC. For this reason, most 14-3-3 in the cell is not perturbed by these drugs. The data from eDRAC experiments provides compelling evidence that the 14-3-3 targeted by PAV-431 comprises only a single-digit percent of the total amount of 14-3-3 present in the cellular extract (see **Figures 5** and **6**). The data from photocrosslinking experiments provides evidence that this targeted subfraction of 14-3-3 is present in an MPC (see **Figures 5** and **6**). PAV-431 was determined to directly bind 14-3-3 but it also was found to indirectly bind multiple other proteins including p62/SQSTM1, VCP, and CAPN2 that are present in the MPC (see **Figures 6A-D**). Since 14-3-3 is known to regulate an array of cellular functions, data showing that PAV-431 targets a particular MPC provides a plausible explanation for why some but not all functions of 14-3-3 are regulated by the compound. The selectivity of PAV-431 to a small subset of 14-3-3 that is specific to a particular MPC or biochemical pathway makes the possibility of developing the chemical series as a 14-3-3 targeting therapeutic increasingly viable.

We originally termed hit compounds identified by our CFPSA screen as ‘assembly modulators’ because they blocked the assembly of viral proteins. However, based on the eDRAC and photocrosslinking results we would propose a more nuanced model for understanding the mechanism of action of assembly modulating compounds. Our data suggests that viral infection modifies a multi-protein complex with catalytic activity to serve multiple alternate needs for the virus. This includes both promoting viral propagation through capsid assembly and blocking innate immune defenses, such as p62/SQSTM1-mediated autophagy (see **Figures 5** and **6**). This model is supported by the changes observed to the MPC targeted by PAV-431 under different conditions which indicate that the target’s composition is dynamic (see **Figure 5**). When cells are infected by viruses, certain proteins appear to be recruited and others appear to be expelled from the targeted multi-protein complex. When infected cells are treated with PAV-431, the reverse happens and the protein composition of the multi-protein complex appears to be largely restored to what was observed in uninfected cells.

The significance of 14-3-3 as the direct binding partner of PAV-431 may be found in its known roles as an allosteric modulator of protein-protein interactions (67) and in its known participation in host anti-viral defenses (**Figure 7B**) (76). This may also, at least in part, account for the antiviral activity observed for PAV-431 against six diverse families of viruses causing human respiratory disease. The drug-binding site within 14-3-3 may represent a ‘high value’ site which multiple viruses have found and exploited over deep evolutionary time. The relationship between particular proteins which comprise the targeted MPC and 14-3-3 as the direct drug-binding partner is unknown besides the evidence that they are transiently co-associated, and the observation that many of these proteins are implicated in the literature as being part of disease-relevant protein-protein interactomes (46–49). While more data is needed, the potential significance of these early results involving the PAV-431 drug target is underscored by the loss upon viral infection, and return upon drug-treatment of infected cells, of p62, a known regulator of autophagy (50,51,66). An inability to trigger innate immune responses after viral infection would be to the virus’s benefit and the host’s detriment. Conversely, restoration of this function would bolster the host’s ability to fend of infection. Thus these compounds appear to have a dual mechanism of action: blockade of viral replication (capsid assembly) and restoration of autophagy, a branch of the innate immune system.

**Figure 7.**
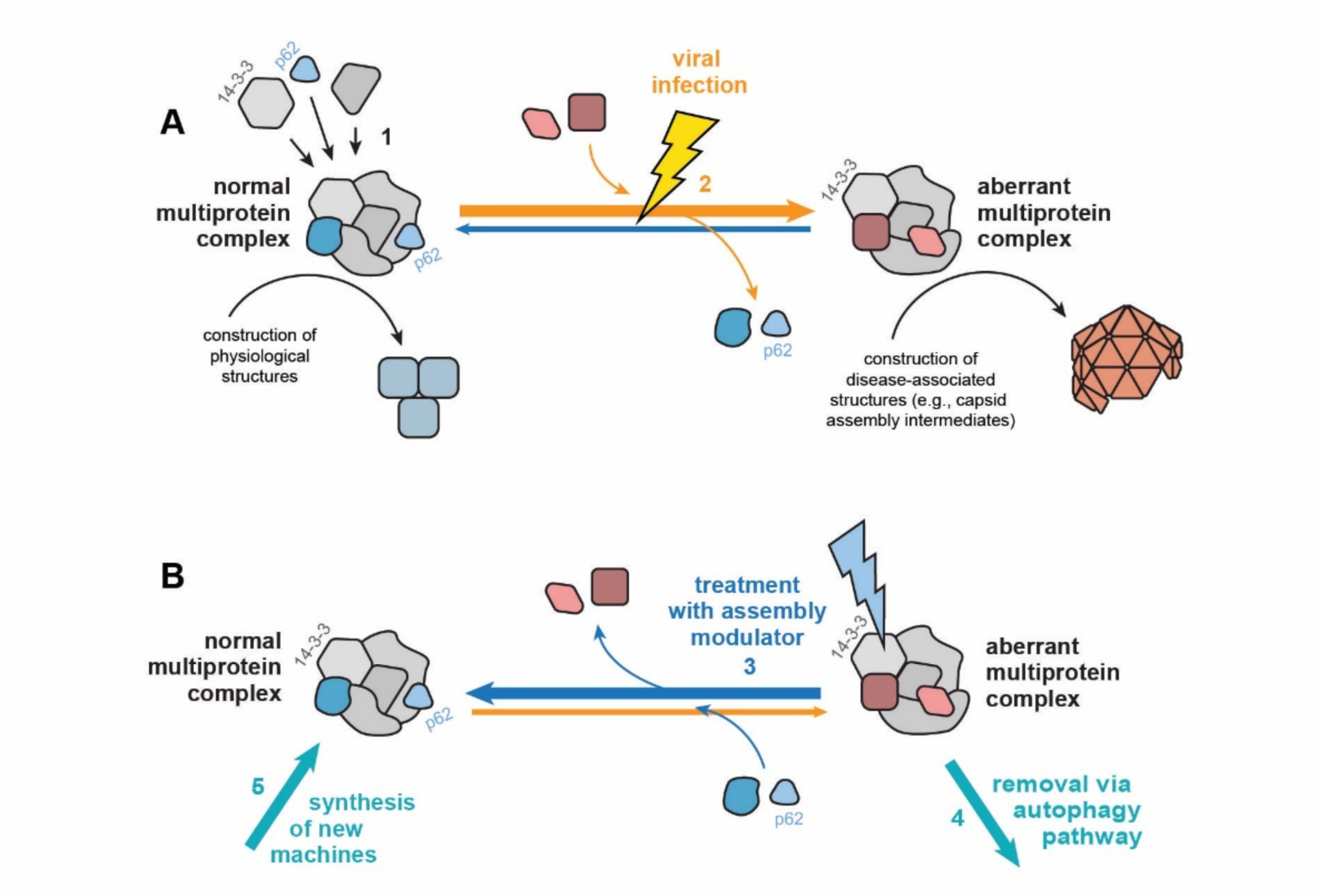
Cartoon diagram of proposed mechanism of action of assembly modulating compounds. **Figure 7 A** illustrates the proposed model where a “normal” MPC with catalytic activity that plays a role in carrying out cellular events in the service of homeostasis is modified to an “aberrant” MPC by a viral infection. The aberrant MPC carries out a reaction which does not serve homeostasis (e.g. building a viral capsid) and perhaps fails to conduct a key event that it should (e.g. inform innate immune mechanisms that the cell is under attack). **Figure 7B** illustrates the proposed mechanism in which treatment with a assembly modulating compound, such as PAV-431, normalizes the complex and its homeostatic functions. The protein 14-3-3 is included in the diagram because it is the protein which the compound appears to directly bind. Its known role as an allosteric regulator may provide insight into how this normalization is achieved.

While the identification of the pan-respiratory assembly modulating chemotype was achieved through unconventional methods, and its novel mechanism of action remains poorly understood, the antiviral activity of compounds from the series have been validated against infectious viruses in both cell culture and animals (see **Figures 2-4**). Cell culture studies, including in primary bronchial epithelial cells cultured at an air-liquid interface and infected with SARS-CoV-2, a model considered as the gold standard for translatability into human therapeutics (77), confirmed antiviral potency of these compounds (see **Figure 4**). Animal studies validated efficacy for survival in an actual pig coronavirus disease and viral load reduction in the cotton rat model of RSV infection (see **Figures 3** and **4**). The path to develop this chemical series to a clinical drug-candidate and conducting IND enabling studies, IND filing, and human clinical trials on the lead compound is straightforward, especially since a more advanced chemical analog displaying substantial improvement in antiviral activity has already been identified (53, 54). If validated in humans, the assembly modulating compounds presented here may have transformative implications for the treatment of respiratory viral disease, applicable to everything from seasonal influenza, common ‘winter viruses’ (HRV, etc), emerging variants of SARS-Cov-2, and any other particularly virulent strains of respiratory disease-causing viruses such as avian influenza.

Future studies with advanced analogs will address the question of whether it is possible to identify analogs that show selectivity for the form of the target observed in viral infection and avoid the form present in the healthy (uninfected) host. Such compounds would be ideal for human therapeutics and predicted to display striking diminution in toxicity.

## Supplemental Figures

**Supplemental Figure 1.**
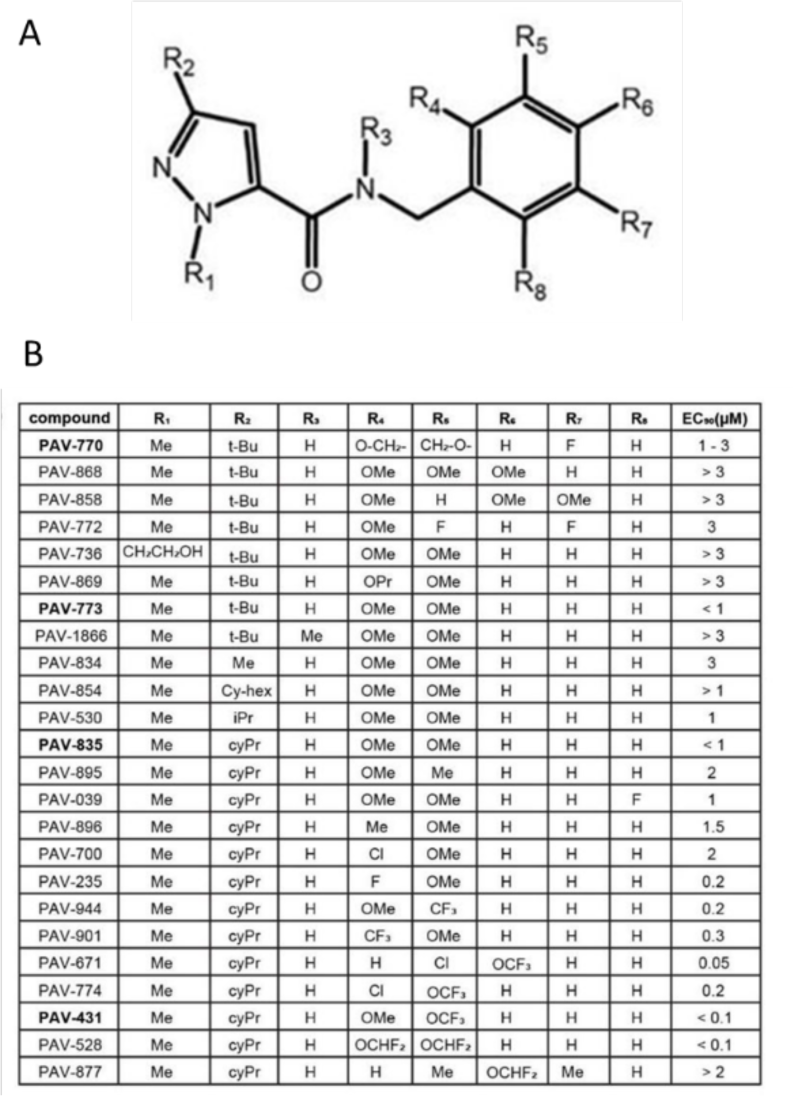
Early SAR exploration within hit chemical series. **Supplemental Figure 1A** shows a Markush structure for the initial hit chemical series in the CFPSA flu capsid assembly screen. **Supplemental Figure 1B** shows the initial structure-activity-relationship pursued to characterize how changes to different parts of the molecule affect activity of the series. EC50 for each compound was determined by TCID_50_ with infectious FLUV in MDCK cells.

**Supplemental Figure 2.**
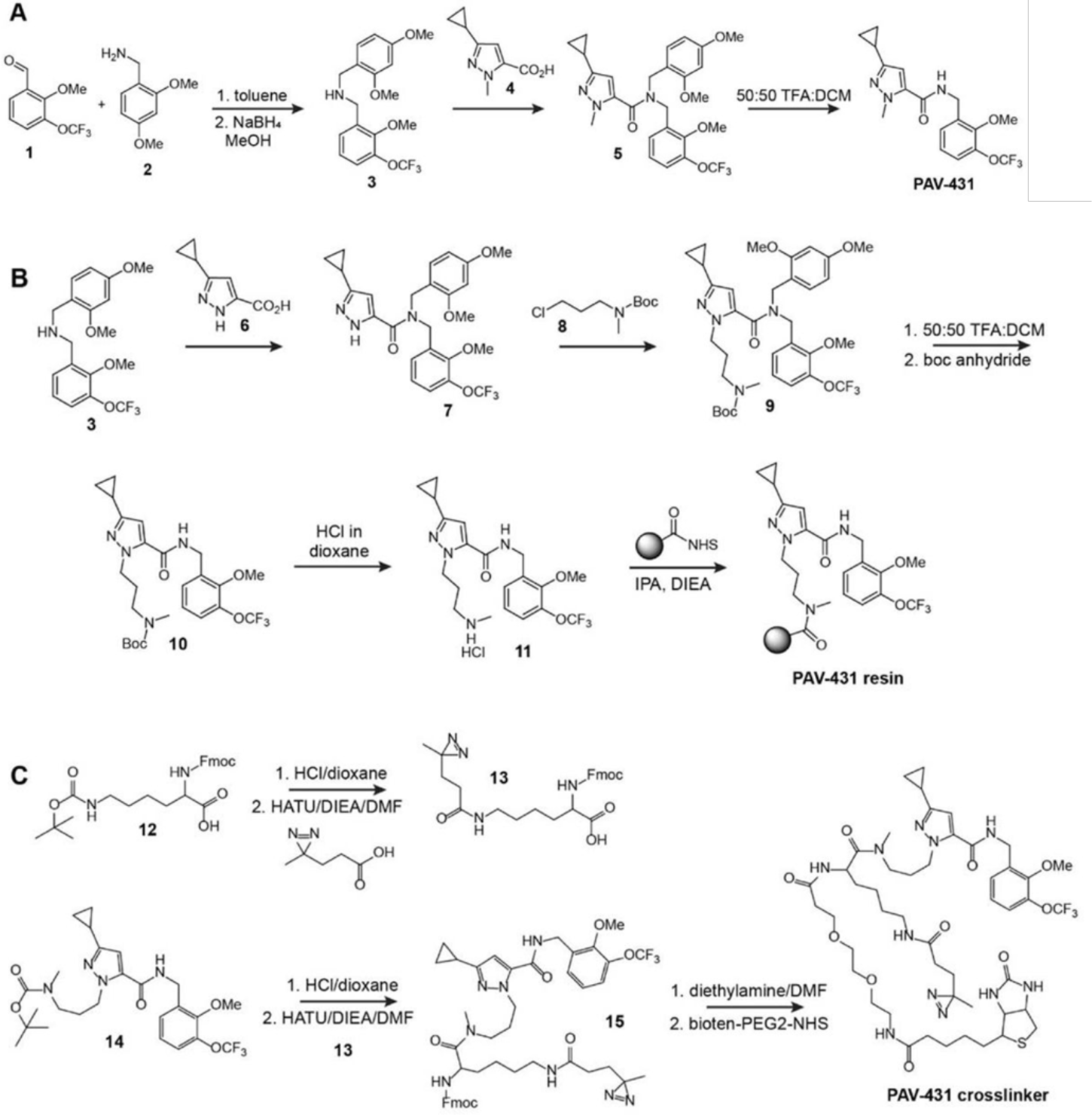
Synthetic scheme for PAV-431 and its resin and photocrosslinker analogs. **Supplemental Figure 2A** shows the synthetic scheme for PAV-431. **Supplemental Figure 2B** shows the synthetic scheme for attachment to a resin by the pyrazole position, which was used in the eDRAC experiments described in Figure 5 and Supplemental Figure 4. The eDRAC experiments described in Figure 6 were conducted with a resin attached from the benzyl ring. **Supplemental Figure 2C** shows the synthetic scheme for the PAV-431 photocrosslinker analog used in the experiments described in Figure 6.

**Supplemental Figure 3.**
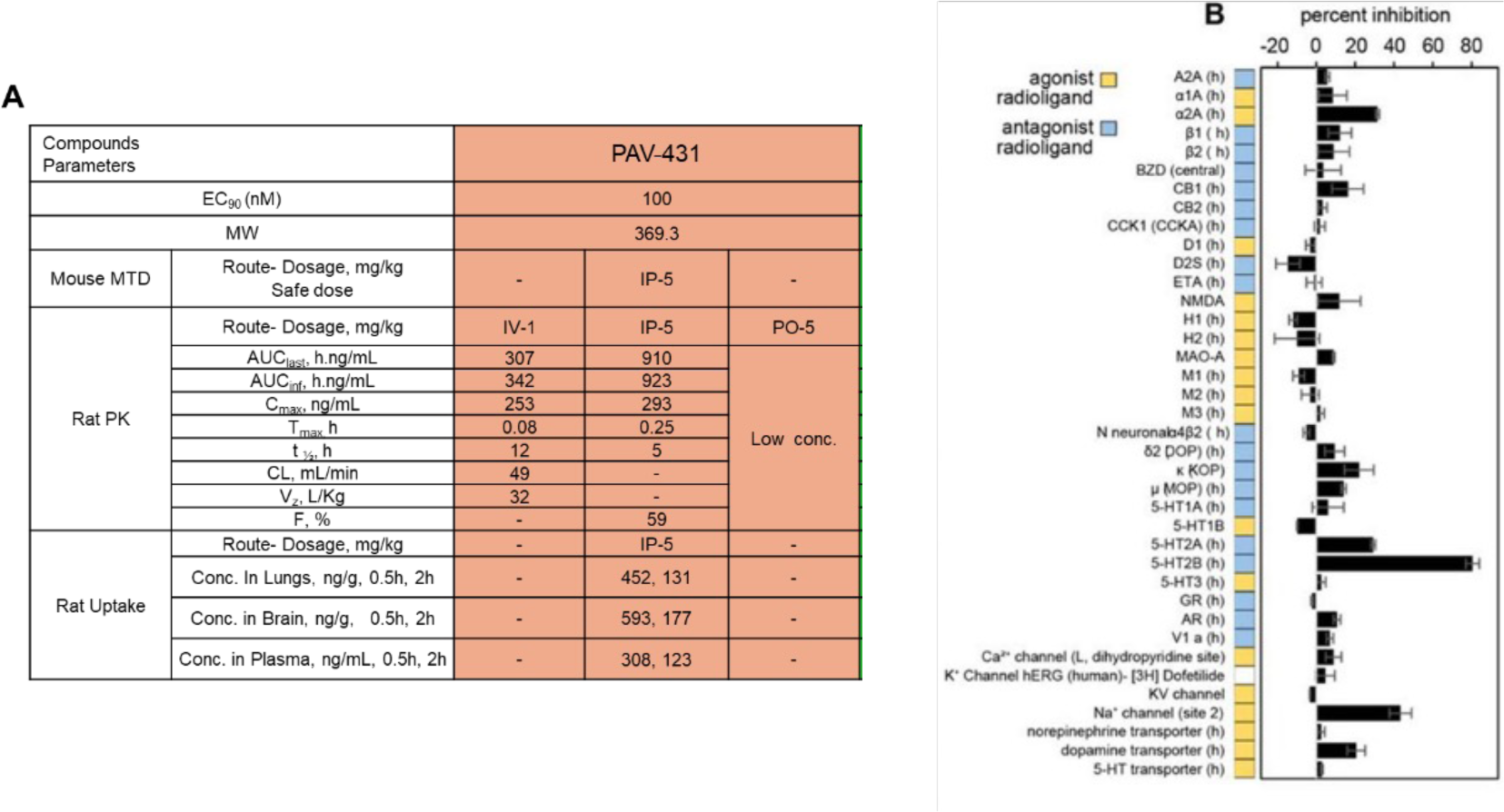
**Supplemental Figure 3A** shows the drug-like properties of PAV-431 including in vivo and in vitro assessments of toxicity as well as pharmacokinetic properties. Maximum tolerated dose (MTD) studies in mice were conducted using female Balb/c mice where randomized groups containing 3 mice were dosed with a single dose of vehicle or compound and monitored for 48 hours for symptoms of toxicity. Pharmacokinetic (PK) studies were conducted in male Sprague Dawley rats where randomixed groups of four animals were administered compound and plasma was collected before dosing then after 5 minutes, 15 minutes, 30 minutes, 1 hour, 2 hours, 4 hours, 8 hours, 12 hours, and 24 hours to determine concentration of the compound in plasma over time. In the uptake studies, animals were euthanized after 30 minutes or 2 hours to determine concentration of the compound in the lung and brain. **Supplemental Figure 3B** shows the results of PAV-431 in an in vitro Cerep panel, a commercial screen for potential to bind to a broad panel of receptors, enzymes, and ion channels, reported as percent inhibition of control specific binding. PAV-431 was tested at 50uM, a concentration ∼500x higher than antiviral EC50. Data shown are the averages of replicates, error bars indicate standard error.

## Materials and methods

### Lead contact and Materials Availability

Further information and requests for resources and reagents should be directed to and will be fulfilled by the Lead Contact Vishwanath R. Lingappa.

Use of unique compound PAV-431 may be available upon request by the Lead Contact if sought for experimental purposes under a valid completed Materials Transfer Agreement.

The number of replicates carried out for each experiment is described in the figure/table legends.

### Chemical Synthesis (see Supplemental Figure 2)

#### Synthesis of PAV-431

Synthetic schemes are illustrated in Figure S6. To a solution of 2-methoxy-3-trifluoromethoxy-benzaldehyde 1 (2.14 g, 9.71 mmol, 1.0 eq) in toluene (20 mL) was added 2,4-dimethoxybenzyl amine 2 (1.78 g, 10.68 mmol, 1.1 eq) and the reaction mixture was stirred at room temperature for 24 hours. Toluene was removed to give a residue, which was taken in MeOH (20 mL) and then NaBH4 (735 mg, 19.42 mmol, 2.0 eq) was added slowly. The reaction mixture was stirred at room temperature for 6 hours. The solvent was removed and the residue was extracted in ethyl acetate and stirred with saturated aq NaHCO3 for 1 hour. The organic layer was collected, dried, and the solvent was removed to give the crude amine 3, which was used in the next step without further purification. To a solution of the crude amine 3 (4.86 mmol, 1.0 eq) in DMF (20 mL) were added the acid 4 (888 mg, 5.35 mmol, 1.1 eq), DIEA (3.13 g, 24.3 mmol, 5eq) and HBTU (2.22 g, 5.83 mmol, 1.2 eq) and the reaction mixture was stirred at room temperature for 12 hours. The reaction mixture was then diluted with ethyl acetate (75 mL) and washed with 10% aq HCl (1 x 50 mL), sat NaHCO3 (1 x 50 mL) and water (4 x 50 mL). The organic layer was collected, dried (MgSO4) and evaporated to give a crude product, which was purified by column chromatography (EtOAc:Hexane 25%:75%)) to give the amide 5, which was directly used in the next step. The amide 5 was treated with 95% TFA:H2O for 12 hours. TFA was removed and azeotroped with toluene to give a residue, which was purified by column chromatography (EtOAc:Hexane 10%:50%) to give PAV-431 (985 mg, > 95% purity).

#### Synthesis of PAV-431 Resin

To a solution of amine 3 (5.85 g, 15.77 mmol, 1.0 eq) in DMF (30 mL) were added the acid 6 (2.38 g, 15.77 mmol, 1.0 eq), DIEA (10.2 g, 78.85 mmol, 5eq) and HBTU (7.17 g, 18.92 mmol, 1.2 eq) and the reaction mixture was stirred at room temperature for 12 hours. The reaction mixture was then diluted with ethyl acetate (75 mL) and washed with 10% aq HCl (1 x 50 mL), sat NaHCO3 (1 x 50 mL) and water (4 x 50 mL). The organic layer was collected, dried (MgSO4) and evaporated to give a crude product, which was purified by column chromatography (EtOAc/Hexane) to give compound 7. To a stirred solution compound 7 (0.8 g, 1.77 mmol, 1.0 eq) and cesium carbonate (1.15 g, 3.54 mmol, 2.0 eq) in DMF (10 mL) was added chloride 8 (0.55 g, 2.66 mmol, 1.5 eq) and the reaction mixture was stirred at room temperature for 24 hours. The reaction mixture was diluted with ethyl acetate and washed with water (4x) and aq NaCl solution. The organic layer was collected, dried (MgSO4) and evaporated to give a crude product, which was purified by column chromatography (EtOAc/Hexane) to give compound 9. The amide 9 (1.0 g, 1.6 mmol) was taken in 95% TFA: H2O and the reaction mixture was for 12 hours. TFA was removed and azeotroped with toluene to give a residue. The residue was taken in DCM and sat. NaHCO3 solution added and stirred for 30 min. The aqueous layer was washed with DCM (2x) and the combined organic layer, dried (MgSO4) and evaporated to give a crude amine, which was used in the next step without purification. To a solution of the crude amine (1.6 mmol, 1.0 eq) and DIEA (412.8 mg, 3.2 mmol, 2.0 eq) in DCM (20 mL), was added boc anhydride (523.2 mg, 2.4 mmol, 1.5 eq) and the reaction mixture was stirred at room temperature for 8 hours. The solvent was removed and the residue was purified by column chromatography (EtOAc/Hexane) to give compound 10. Compound 10 (100 mg, 0.19 mmol) was in 5 mL of DCM and then 4 M HCl in dioxane (3 mL, 12 mmol) was added and the reaction mixture was stirred for 12 hours. Solvents were removed to give compound 11 as a HCl salt, which was used in the next step without further purification. To a solution of Affi-Gel 10 (Bio-Rad, 2 ml, 0.03 mmol, 1.0 eq) in a solid phase synthesis tube with frit was added a solution of compound 11 (27.7 mg, 0.06 mmol, 2.0 eq) and DIEA (1.0 mL) in isopropyl alcohol (4 mL) and the tube was put in a shaker for 12 hours. Excess reagents were drained and the resin was washed with isopropyl alcohol (3x) and then saved in isopropyl alcohol.

#### Synthesis of PAV-431 Photocrosslinker

To 6-(tert-Butoxycarbonylamino)-2-(9H-fluoren-9-ylmethoxycarbonylamino)hexanoic acid 12 [468mg (1mmol)] in a 40ml screw top vial was added 4N HCl in Dioxane (3ml). The vial was sealed and gently agitated for 20 minutes at room temperature. The mix was then rotary evaporated to dryness and the residue placed under high vacuum overnight. The dried residue was taken up into 4ml of DMF (anhydrous) and then sequentially treated with 3-(3-Methyldiazirin-3-yl)propanoic acid [128mg (1mmol)](42), and DIEA [695ul (4mmol)]. With rapid stirring, under Argon atmosphere, was added dropwise HATU [380mg (1mmol)] dissolved in 1ml of DMF. After stirring for 30 minutes the mixture was quenched with 10ml of sat. NH4Cl solution and then extracted 2 x with 10ml of EtOAc. The combined organic extracts were washed once with sat. NaCl, dried (Mg2SO4) and then rotary evaporated to dryness. The residue was purified by flash chromatography, using a gradient of Ethyl acetate and Hexane, affording 2-(9H-fluoren-9-ylmethoxycarbonylamino)-6-[3-(3-methyldiazirin-3-yl)propanoylamino]hexanoic acid 13 (293mg) in 61% yield.

To tert-Butyl N-[3-[3-cyclopropyl-5-[[2-methoxy-3-(trifluoromethoxy)phenyl]methylcarbamoyl]pyrazol-1-yl]propyl]-N-methyl-carbamate 14 [16mg (0.03 mmol)] in a 40ml screw top vial was added 4N HCl in Dioxane (0.5ml). The vial was sealed and gently agitated for 20min at room temperature. The mix was then rotary evaporated to dryness and the residue placed on high vacuum overnight. The dried residue was taken up into 1ml of DMF (anhydrous) and then sequentially treated with compound 13 [14.5mg (0.03mmol)], and DIEA [32ul (0.18mmol)]. With rapid stirring, under Argon atmosphere, was added dropwise HATU [14.6mg (0.038mmol)] dissolved in 300ul of DMF. After stirring for 30 min the mixture was quenched with 5ml of sat. NH4Cl solution and then extracted 2 x with 5ml of EtOAc.

The combined organic extracts were washed once with sat. NaCl, dried (Mg2SO4) and then rotary evaporated to dryness. The residue was purified by flash chromatography, using a gradient of Ethyl acetate and Hexane, affording 9H-fluoren-9-ylmethyl N-[1-[3-[3-cyclopropyl-5-[[2-methoxy-3-(trifluoromethoxy)phenyl]methylcarbamoyl]pyrazol-1-yl]propyl-methyl-carbamoyl]-5-[3-(3-methyldiazirin-3-yl)propanoylamino]pentyl]carbamate 15 (28mg) in quantitative yield.

To compound 15 [28mg (0.03 mmol)] in a 40ml screw top vial was added 50/50 Diethylamine / DMF (0.5ml). The vial was sealed and gently agitated for 60min at room temperature. The mix was then rotary evaporated to dryness and the residue placed on high vacuum overnight. The residue was triturated 2 x with 3ml of Hexane to remove the Dibenzofulvene amine adduct. The residue was again briefly placed on high vacuum to remove traces of Hexane. The dried residue was taken up into 1ml of DMF (anhydrous) and then treated with Biotin-PEG2-NHS [15mg (0.03mmol)] (purchased from ChemPep), and DIEA [16ul (0.09mmol)] and then purged with Argon. After stirring overnight at room temperature, the mixture was rotary evaporated to dryness. The residue was purified by reverse phase prep chromatography, using a gradient of 0.1% TFA water and Acetonitrile, affording 5-cyclopropyl-N-[[2-methoxy-3-(trifluoromethoxy)phenyl]methyl]-2-[3-[methyl-[6-[3-(3-methyldiazirin-3-yl)propanoylamino]-2-[3-[2-[2-[5-(2-oxo-1,3,3a,4,6,6a-hexahydrothieno[3,4-d]imidazol-4-yl)pentanoylamino]ethoxy]ethoxy]propanoylamino]hexanoyl]amino]propyl]pyrazole-3-carboxamide (26mg) in 80% yield. All compounds were confirmed by LCMS.

## Method and Analysis Details

### *In vitro* studies

#### CFPSA screen

Coding regions of interest were engineered behind the SP6 bacteriophage promoter and the Xenopus globin 5ʹ UTR63. DNA was amplified by PCR and then transcribed in vitro to generate mRNA encoding each full-length protein. Translations were carried out in wheat germ extracts supplemented with energy and amino acids, as previously described(7). Moderate-throughput small molecule screening was carried out in 384-well plate format by translation of eGFP and FLUV NP and M mRNA in the presence of small molecules from the Prosetta compound collection (Figure S2). Reactions were run at 26°C for 1-2 hours for synthesis, followed by assembly at 34°C for 2 hours. eGFP fluorescent readout was measured at 488/515 nm (excitation/emission) to assess protein synthesis. Assembly products were captured on a second 384-well plate precoated with affinity-purified FLUV NP antibody. Plates were washed with PBS containing 1% Triton X-100, decorated with biotinylated affinity-purified FLUV NP antibody, washed, detected by NeutraAvidin HRP, washed again, and then incubated with a fluorogenic HRP substrate Quanta Blue for 1 hour. FLUV assembly fluorescent readout was measured at 330/425 nm (excitation/emission).

#### FLUV assay in MDCK cells

MDCK.2 cells were seeded at 3×104 cells/well in Eagle’s minimal essential medium (MEM) supplemented with fetal bovine serum (FBS) in a 96-well plate and incubated overnight at 37°C. The next day, cells were washed with phosphate buffered saline (PBS) and infected with FLUV A/WSN/33 at an MOI of 0.01-0.001 for 1 hour, after which the virus containing media was removed and fresh media containing dilutions of compound or DMSO as a vehicle control was added to the cells. After 24 hours, media was removed, cells were washed with PBS, and fresh media was added for a 2 hour incubation and then collected for TCID50 determination. Seven replicates of 10-fold serial dilutions of collected media were added to new cells and incubated at 37°C for 3 days. The number of infected wells for each dilution was determined by visual inspection, and TCID50/mL was calculated using the Reed and Muench method. Infection experiments were conducted in a BSL2 laboratory.

#### BoCoV assay in HRT-18G cells

HRT-18G cells were seeded at 3×104 cells/well in Dulbecco’s modified Eagle medium (DMEM) in a 96-well plate and incubated overnight at 37°C. The next day, cells were infected with BoCoV BRCV-OK-0514-2 (ATCC VR-2460) at an MOI of 1 for 2 hours, after which the virus containing media was removed, cells were washed with PBS, and fresh media containing dilutions of compound or DMSO as a vehicle control was added to the cells. After 42-48 hours, media was removed, cells were washed with PBS, and fresh media was added for a 4 hour incubation and then collected for TCID50 determination. Infection experiments were conducted in a BSL2 laboratory.

#### HRV assay in H1-HeLa cells

H1-HeLa cells were seeded at 7×104 cells/well in MEM in a 96-well plate and incubated overnight at 37°C. The next day, cells were infected with HRV-16 at an MOI of 5 for 1.5 hours, after which the virus containing media was removed, cells were washed with PBS, and fresh media containing dilutions of compound or DMSO as a vehicle control was added to the cells. After 72 hours, media was collected for TCID50 determination. Infection experiments were conducted in a BSL2 laboratory.

#### MHV assay in BHK-21 cells

BHK-21 cells were seeded at 2.5×105 cells/well in MEM in a 96-well plate and incubated overnight at 37°C. The next day, cells were infected with MHV-68 at an MOI of 0.5 for 1.5-2 hours, after which the virus containing media was removed, cells were washed with PBS, and fresh media containing dilutions of compound or DMSO as a vehicle control was added to the cells. After 24 hours, media was removed, cells were washed with PBS, and fresh media was added for a 4 hour incubation and then collected for TCID50 determination. Infection experiments were conducted in a BSL2 laboratory.

#### SARS-CoV-2 assay in Vero cells

Vero clone E6 (CRL-1586) cells were plated at 3×105 cells/well in DMEM in 6-well plates and incubated overnight at 37°C. The next day, cells were washed once with PBS and then infected with SARS-CoV-2 WA1/2020 (MN985325.1, BEI resources) at a MOI of 0.01 for 1 hour after which the virus containing media was removed and the compounds were added to the cells and incubated for 72 hours at 37°C at 5% CO2. The cells were then fixed and stained with crystal violet to determine plaque numbers(38). Infection experiments were conducted in a BSL3 laboratory. Data shown in Figure 4B are the averages of two biological replicates; error bars indicate standard error; DMSO is included as the vehicle control.

#### SARS-CoV-2 (delta) assay in Calu-3 cells

Calu-3 cells were seeded at a density of 3×104 cells/well in DMEM in 96-well plates and incubated overnight at 37°C. The next day, cells were pre-incubated with compounds for 4 hours before they were infected with SARS-CoV-2 delta SL102 (EPI_ISL_4471559) at a MOI of 0.01-0.05. After 24 hours the viruses within 50 µl of the supernatants were lysed with 200 µL AVL-buffer (Qiagen) and 200 µL 100% ethanol was added for complete inactivation. RNA was extracted from 200 µL of the lysates using the EZ1 Virus Mini-Kit (Qiagen), and analyzed by qPCR as described(39). Infection experiments were conducted in a BSL3 laboratory. Data shown are the averages of three biological replicates; error bars indicate standard error; DMSO is included as the vehicle control.

#### Recombinant ZsGreen-expressing Nipah virus infection

HSAEC1-KT cells were seeded at 10,000 cells per well the day prior to infection in 96-well black plates with clear bottoms (Costar 3603). The following day, cells were infected with recombinant Nipah virus expressing ZsGreen fluorescence protein (rNiV-ZsG) (Lo et al., 2014, 2018, 2020 AVR: Welch et al., 2020 JID) at multiplicity of infection 0.01 with ∼ 100 50% tissue culture infectious dose (TCID50). Levels of rNiV-ZsG replication were measured at 72 hour post-infection based on mean ZsGreen fluorescence signal intensity (418ex/518em) using a Biotek HD1 Synergy instrument (Aglilent). Fluorescence signal intensity assayed in DMSO-treated, virus-infected cells were set as 100% ZsGreen fluorescence. Data points and error bars for all reporter assays indicate the mean value and standard deviation of 4 biological replicates, and are representative of at least 2 independent experiments in HSAEC1-KT cells. Concentrations of compound that inhibited 50% of the green fluorescence signal (EC50) were calculated from dose response data fitted to the mean value of experiments performed for each concentration in the 10-point, 3-fold dilution series using a 4-parameter non-linear logistic regression curve with variable slope using GraphPad Prism 9 (GraphPad Software, La Jolla, CA, USA).

#### CellTiterGlo cell viability assay

Cell viability was assayed using CellTiter-Glo 2.0 assay reagent (Promega) according to manufacturer’s recommendations, with luminescence measured at 72 hours post-compound treatment using a Biotek HD1 Synergy instrument. Luminescence levels (indicative of cellular ATP levels as a surrogate marker of cell viability) assayed in DMSO-treated, uninfected cells were set as 100% cell viability. Dose response curves were fitted to the mean value of experiments performed for each concentration in the 10-point, 3-fold dilution series using a 4-parameter non-linear logistic regression curve with variable slope. All CellTiter-Glo cell viability assays were conducted in 96-well opaque white plates (Costar 3917). Concentrations of compound that inhibited 50% of the luminescence signal (CC50) were calculated from dose response data fitted to the mean value of experiments performed for each concentration in the 10-point, 3-fold dilution series using a 4-parameter non-linear logistic regression curve with variable slope using GraphPad Prism 9 (GraphPad Software, La Jolla, CA, USA).

#### Primary airway epithelial cell culture

Human bronchus was harvested from 3 explanted lungs. The tissue was submerged and agitated for 1 minute in PBS with antibiotics and 5mM dithiothreitol to wash and remove mucus. After 3 washes, the tissue was placed in DMEM with 0.1% protease and antibiotics overnight at 4°C. The next day the solution was agitated and remaining tissue removed. Cells were centrifuged at 300g/4°C for 5 minutes, then resuspended in 0.05% trypsin-EDTA and incubated for 5 minutes at 37°C. The trypsinization reaction was neutralized with 10% FBS in DMEM, then cells were filtered through a cell strainer and centrifuged at 300g/4°C for 5 minutes. The cell pellet was resuspended in 10% FBS in DMEM and a 10uL aliquot was stained with trypan-blue and counted on a hemocytometer. 7.5×104 cells were plated onto each 6mm/0.4mm FNC-coated Transwell air-liquid interface (ALI) insert. 10% FBS in DMEM and ALI media were added in equal volumes to each basal compartment and cultures were incubated at 37°C/5% CO2. The next day, media was removed and both compartments were washed with PBS and antibiotics. ALI media was then added to each basal compartment and changed every 3 days until cells were ready for use at day 28.

All studies involving SARS-CoV-2 infection of primary airway epithelial cells were conducted in the Vitalant Research Institute BSL3 High-Containment Facility. 6 hours prior to infection, ALI medium containing dilutions of drugs (100nM) or DMSO was added to the basal compartment. For infection, ALI medium containing drugs was removed, and SARS-CoV-2 diluted in ALI-culture medium containing drugs (100nM, MOI=0.1) was added on to the apical chamber of inserts (250 µl) and the basal compartment (500 µl). The cultures were incubated for 2 hours at 37℃/5% CO2 to allow for virus entry, then washed, and 500 µl of fresh ALI medium containing drugs (100 nM) was added to the basal compartment. Drugs were maintained in the medium for the duration of the experiment. Cells were incubated at 37℃/5% CO2 and harvested for analysis at 36 hours post-infection.

Total RNA was extracted from mock and SARS-CoV-2-infected primary airway epithelial cells with or without drug treatment lysed in Trizol (Thermo Fisher Scientific) using the chloroform-isopropanol-ethanol method. 500 ng of RNA was reversed transcribed into cDNA in 20 uL reaction volume using RevertAid First Strand cDNA Synthesis kit (Thermo Fisher) in accordance to the manufacturer’s guidelines. RT-PCR was performed for each sample using TaqmanTM Universal Master Mix II, with UNG (Thermo Fisher) on the ViiA7 Real time PCR system. Primers and probes (2019-nCoV RUO kit) for detection of the SARS-CoV-2 Nucleocapsid (N) gene were obtained from IDT.

#### Alamar Blue HS cell viability assay

Cell viability was assayed using Alamar Blue HS reagent (Thermofisher) according to manufacturer’s recommendations, with fluorescence (560ex/590em) measured at 72 hours post-compound treatment after 4 hours of incubation with reagent using a Biotek HD1 Synergy instrument. Fluorescence levels (indicative of resazurin reduction as a surrogate marker of cell viability) assayed in DMSO-treated, uninfected cells were set as 100% cell viability. Dose response curves were fitted to the mean value of experiments performed for each concentration in the 10-point, 3-fold dilution series using a 4-parameter non-linear logistic regression curve with variable slope. All Alamar Blue assays were conducted in 96-well black plates with clear bottoms. Concentrations of compound that inhibited 50% of the fluorescence signal (CC50) were calculated from dose response data fitted to the mean value of experiments performed for each concentration in the 10-point, 3-fold dilution series using a 4-parameter non-linear logistic regression curve with variable slope using GraphPad Prism 9 (GraphPad Software, La Jolla, CA, USA).

### Cell lysate preparation

Cells or tissues were extracted with PB buffer (10 mM Tris pH 7.6, 10 mM NaCl, 0.1 mM EDTA, and 0.35% Triton X-100), and centrifuged at 10,000 x g for 10 min. The supernatants were collected and flash frozen.

### Energy-dependent drug resin affinity chromatography (eDRAC)

Drug resin was prepared by coupling compound PAV-431 to an Affi-gel resin at a concentration of 10 µM via the pyrazole nitrogen (Figure S6, synthetic chemistry described below), or position 4 of the phenyl group. Control resin was prepared by blocking the Affi-gel matrix without drug. Resins were equilibrated with column buffer (50 mM HEPES, pH 7.6, 100 mM KAc, 6 mM MgAc, 1 mM EDTA, 4 mM TGA) prior to any DRAC experiments. 30 µL of cell extract supplemented with energy (1 mM ATP, GTP, CTP and UTP with 4 mM creatine phosphate, and in some cases 5 µg/ml rabbit creatine kinase) was applied to resin columns. The columns were clamped and incubated at 22°C for 1 hour for binding, and flow through was collected. The columns were then washed with 100 bed volumes of column buffer. For elution of bound complexes, 100 µL of column buffer containing free drug at a final concentration of 100 µM – 1 mM (approaching its maximum solubility in water) and supplemented with energy was added, the column was clamped for 1 hour, and serial eluates were collected. Eluates were analyzed by SDS-PAGE and WB. or later use.

### Western blotting

SDS-PAGE gels were transferred in Towbin buffer to a polyvinylidene fluoride membrane. Membranes were then blocked in 1% BSA, incubated for 1 hour at room temperature in a 1:1000 dilution of 100 μg/mL affinity-purified primary antibody, washed three times in PBS with 0.1% Tween-20, incubated for 1 hour in a 1:5000 dilution of secondary anti-rabbit or anti-mouse antibody coupled to alkaline phosphatase, washed further, and incubated in developer solution prepared from 100 μL of 7.5 mg/mL 5-bromo-4-chloro-3-indolyl phosphate dissolved in 60% dimethyl formamide (DMF) in water and 100 μL of 15 mg/mL nitro blue tetrazolium dissolved in 70% DMF in water, adjusted to 50 mL with 0.1 M Tris (pH 9.5)/0.1 mM magnesium chloride.

### MS-MS analysis

Samples were processed by SDS-PAGE using a 10% Bis-Tris NuPAGE gel (Invitrogen) with the MES buffer system. The mobility region was excised and processed by in-gel digestion with trypsin using a ProGest robot (Digilab) with the protocol outlined below. Washed with 25 mM ammonium bicarbonate followed by acetonitrile. Reduced with 10 mM dithiothreitol at 60°C followed by alkylation with 50 mM iodoacetamide at room temperature. Digested with trypsin (Promega) at 37°C for 4 hours. Quenched with formic acid, lyophilized, and reconstituted in 0.1% trifluoroacetic acid.

Half of each digested sample was analyzed by nano LC-MS/MS with a Waters M-Class HPLC system interfaced to a ThermoFisher Fusion Lumos mass spectrometer. Peptides were loaded on a trapping column and eluted over a 75 μm analytical column at 350 nL/min; both columns were packed with Luna C18 resin (Phenomenex). The mass spectrometer was operated in data-dependent mode, with the Orbitrap operating at 60,000 FWHM and 15,000 FWHM for MS and MS/MS respectively. APD was enabled and the instrument was run with a 3 s cycle for MS and MS/MS.

Data were searched using a local copy of Mascot (Matrix Science) with the following parameters: Enzyme: Trypsin/P; Database: SwissProt Human plus the custom sequences* (concatenated forward and reverse plus common contaminants); Fixed modification: Carbamidomethyl (C)Variable modifications: Oxidation (M), Acetyl (N-term), Pyro-Glu (N-term Q), Deamidation (N/Q)Mass values: Monoisotopic; Peptide Mass Tolerance: 10 ppm; Fragment Mass Tolerance: 0.02 Da; Max Missed Cleavages: 2. The data was analyzed by label free quantitation (LFQ) methods(40). LFQ intensity values of each condition were measured in triplicate and compared against each other to generate log2 fold change values for each protein and each combination of conditions. Proteins that were found significantly enriched by a log2 fold change of > 1 and an adjusted p-value (accounting for multiple hypothesis testing) of < 0.05 in the FLUV infected eDRAC eluates compared to the uninfected eluates were searched for in a list of high confidence FLUV virus-host protein interactions and the VirusMentha database of virus-protein interactions (46,47). Likewise, significantly enriched and depleted proteins found in the BoCoV infected eDRAC eluate were searched for in a list of high confidence coronavirus interactors and an aggregated list of coronavirus protein interactors shown experimentally (48,49).

### Photocrosslinking and streptavidin precipitation

eDRAC columns were eluted with 100µM PAV-431 photocross-linker at 22oC. Eluates were crosslinked by exposure to UV light for 3 minutes. Crosslinked products were subjected to treatments that maintained protein-protein associations (native) or which reduced and denatured all proteins (denatured). Native conditions were maintained by diluting an aliquot of the product 20x with 1% Triton-X-100 column buffer. Denaturation was achieved by adjusting an aliquot to 1% SDS and 10mM DTT and heating to 100oC/10 minutes prior to 20x dilution with 1% Triton-X-100 column buffer. Streptavidin Sepharose beads were added to both native and denatured samples and mixed for 1 hr to capture all biotinylated proteins, with and without co-associated proteins in the native and denatured cases respectively, then washed 3x with 1% Triton-containing column buffer. Washed beads were resuspended in 20µl of SDS loading buffer and analyzed by SDS-PAGE and WB.

### In vivo studies

#### PEDV pig study

18 litters comprised of 91 individuals of newborn (2 – 4 days old) crossbred pigs weighing 3 kg were randomized to control (vehicle) or treatment groups. Animals were infected with 1×105 PFU of PEDV administered orally. Vehicle or drug was administered intramuscular at 4 mg/kg immediately after challenge and again 24 hours post-infection. Compound efficacy was determined by survivability. Endpoint of study was 6 days post-infection.

#### RSV cotton rat study

Female cotton rats, ∼5 weeks of age, were obtained from Envigo (formerly Harlan), ear-tagged for identification purposes, and allowed to acclimate for > 1 week prior to study start. Animals were housed individually. Vehicle or drug was administered by an intraperitoneal route twice daily on study days −1 through day 4. On day 0, animals were infected with 1×105 PFU of RSV A-2 virus originally obtained from ATCC (VR-1540), administered in a 50 mL volume by an intranasal route approximately 2 hours after the morning treatment dose. Back titration of the viral stock and diluted inoculum was performed to confirm the titer of the RSV stock used for infection. All inoculations were performed while the animals were under the influence of inhalant anesthesia. All animals were euthanized on day 5 and the lungs were processed for determination of RSV titers by plaque assay.

## Acknowledgements

We thank Alfredo Calayag, Lisa Tucker, Caleb Declouette, Yvonne Dickschen, and Björn Wefers for excellent technical assistance, David Hanzel and Homer Boushey for careful reading and improvement of the manuscript, and Dmitry Temnikov for IT support. We are indebted to the late Guenter Blobel for advice, inspiration, and encouragement.

